# Transferability of Brain decoding using Graph Convolutional Networks

**DOI:** 10.1101/2020.06.21.163964

**Authors:** Yu Zhang, Pierre Bellec

## Abstract

Transfer learning has been a very active research topic in natural image processing. But few studies have reported notable benefits of transfer learning on medical imaging. In this study, we sought to investigate the transferability of deep artificial neural networks (DNN) in brain decoding, i.e. inferring brain state using fMRI brain response over a short window. Instead of using pretrained models from ImageNet, we trained our base model on a large-scale neuroimaging dataset using graph convolutional networks (GCN). The transferability of learned graph representations were evaluated under different circumstances, including knowledge transfer across cognitive domains, between different groups of subjects, and among different sites using distinct scanning sequences. We observed a significant performance boost via transfer learning either from the same cognitive domain or from other task domains. But the transferability was highly impacted by the scanner site effect. Specifically, for datasets acquired from the same site using the same scanning sequences, using transferred features highly improved the decoding performance. By contrast, the transferability of representations highly decreased between different sites, with the performance boost reducing from 20% down to 7% for the Motor task and decreasing from 15% to 5% for Working-memory tasks. Our results indicate that in contrast to natural images, the scanning condition, instead of task domain, has a larger impact on feature transfer for medical imaging. With other advanced tools such as layer-wise fine-tuning, the decoding performance can be further improved through learning more site-specific high-level features while retaining the transferred low-level representations of brain dynamics.

## 1 Introduction

Transfer learning has the potential to allow us to train complex models even in the absence of extensive training data, which is highly beneficial in the field of medical imaging, where the lack of a sufficiently large dataset for specific experimental questions is pervasive. Current literature on transfer learning for medical imaging is still dominated by transferring features learned on ImageNet followed by fine-tuning the model on medical images (13; 10). There are however substantial differences between natural and medical images. For instance Gabor filters were widely used for edge detection in natural images but were not detected in medical images (10). The knowledge transfer from natural images to medical images may thus suffer from fundamental limitations (13).

In this study, we specifically focused on the field of decoding brain cognitive states from recorded neural activities using functional magnetic resonance imaging (fMRI). Convergent evidence has shown that neural dynamics are integrated across multiple functional systems, and even the whole brain in order to accomplish a particular cognitive task (11). In this case, the issue of low transferability of features learned from natural images is severe. Therefore, instead of transferring from natural images, we directly trained our base model on a large-scale neuroimaging dataset collected from the Human Connectome Project database. A deep graph neural network consisting of six graph convolutional layers (feature extractor) followed by two fully connected layers (classifier) was used for brain decoding (16). The transferability of brain decoding was evaluated under different circumstances including knowledge transfer across different cognitive domains, different groups of subjects, and even between different sites using different scanning sequences. We also investigate other factors that might have an impact on the transfer learning, including the sample-size of the dataset, the transferability of different layers, and whether to use fixed features or fine-tuning each GCN layer. The interpretability of transfer learning was also investigated by analysing the similarity of learned representations not only between different categories but also between different decoding models, including the model trained from random initialization, transferring features from the base model and after layer-wise fine-tuning of GCN layers. Moreover, in order to evaluate whether the decoding inference was based on biologically meaningful features, we also generated saliency maps for each type of the decoding model and specifically investigated how the saliency evolved after transfer learning and fine-tuning.

## 2 Methods and Materials

### 2.1 fMRI Datasets and Preprocessing

We used the block-design task fMRI data from the Human Connectome Project (HCP) S1200 release (https://db.humanconnectome.org/data/projects/HCP_1200). The minimal preprocessed fMRI data of the CIFTI format were used, which maps individual fMRI time-series onto the standard surface template with 32k vertices per hemisphere. The task fMRI data includes seven cognitive tasks, which are emotion, gambling, language, motor, relational, social, and working memory. Here, we mainly focused on two cognitive tasks: 1) Motor task: participants performed five different types of body movements in scanner, i.e. movement of left and right foot, left and right hand, and tongue, with each task event lasting for 12s; 2) Working-memory task, that involves two-levels of cognitive functions, with a combination of the category recognition task and N-Back memory task. Specifically, participants are presented with pictures of places, tools, faces and body parts in separate blocks, with half of the blocks using a 2-back working memory task (showing the same image after two image blocks) and the other half using a 0-back working memory task (requiring to recognize a single image for the entire duration of one block), each block lasting for 25s. Further details on fMRI data acquisition, task design and preprocessing can be found in (1; 4).

We also used a second task-fMRI dataset acquired from the Individual Brain Charting (IBC) project (9) (https://www.openfmri.org/dataset/ds000244), consisting of 12 subjects performing the same types of cognitive tasks as the HCP database. Modifications have been made in both scanning environments and task protocols, for instance, 1) using a Siemens 3T Magnetom Prismafit scanner at the NeuroSpin platform instead of using the 3T Connectome Scanner at the HCP informatics platform (14); 2) translating all instructions and stimuli presented in experimental paradigms from English to French; 3) using longer acquisition time (TR=2s in IBC, 0.72s in HCP) and more event blocks per run (20 vs 10 blocks for the Motor task, 16 vs 8 blocks for Working-memory tasks). In order to compensate for their difference in TR which might result in capturing different dynamics of task-evoked hemodynamic response, we downsampled the temporal resolution for all HCP datasets by using a time step of three TRs (equivalent TR=2.1s).

### 2.2 Brain State Annotation pipeline

We used the brain state annotation model (16), that has 6 graph convolutional layers with 32 graph filters at each layer followed by 2 fully connected layers as the classifier. The model takes a short series of fMRI data as input, propagates information among inter-connected brain regions and networks, generates a high-order graph representation and finally predicts the corresponding cognitive labels. Specifically, all fMRI volumes were first mapped onto the 360 regions of the Glasser atlas (3), by averaging the BOLD signals within each parcel. Then, the time-series for each event task was extracted, by 1) realigning fMRI volumes with the experimental designs using task onsets and durations; 2) cutting the time-series into bins of the selected time window using a sliding-window approach; 3) downsamping the time-series at the given time step; 4) tag the task labels for each segment of fMRI time-series. The datasets were split into training (70%), validation (10%), test (20%) sets using a subject-specific split scheme, such that time windows collected on the same subject could not appear simultaneously in the training and validation (or testing) set. The base models were trained for 100 epochs with the batch size set to 10 subjects and learning rate set to 0.001, while the other models were trained with a smaller batch size (2 subjects) and learning rate (0.0001) with 150 epochs. The best model with the highest prediction score on the validation set was saved, and then evaluated separately on the test set. We also used a L2 regularization of 0.0005 on weights and a dropout rate of 0.5 on all layers. The implementation of the GCN model for brain decoding was based on Pytorch-1.0, and will be made publicly available.

### 2.3 Transfer learning of brain decoding

We trained the base model for brain decoding using fMRI data from 1000 subjects collected from the HCP database. We chose different time window for the Motor and Working-memory task in order to achieve their peak performance in brain decoding (16), i.e. a 10s window for Motor task (5 fMRI volumes as input) and a 20s window for Working-memory (10 fMRI volumes as input). Two domain-general decoding models were also trained using the above time windows. For task events with a shorter duration than the chosen time window, a neighborhood wrapping method was applied. Here, we used two types of transfer learning, either transferring features from a domain-specific base model trained by exclusively using fMRI signals from the corresponding task domain, or using a domain-general base model trained using fMRI data from all available cognitive tasks. A layer-wise fine-tuning approach was used that gradually tuned the last GCN layers while keeping the first few layers fixed. Another type of transfer learning named transfusion (10) was also evaluated, which only transferring the first few GCN layers while training the rest of network from random initialization. A summary of all models used in this study can be found in Table 1.

**Table 1:** Models for brain decoding and transfer learning.

| Model name<br>abbrev. | Description |
| --- | --- |
| Domain-specific | Training models using fMRI data of one specific task domain<br>e.g. Motor and Working-memory |
| Domain-general | Training models using all available fMRI data under 21 task conditions |
| Base | Pretrained decoding model using 1000 subjects from HCP database |
| Scratch | Training models from random initialization |
| Transfer | Copying and fixing all GCN layers from the base model |
| Freeze $k$ | Freezing the weights up to $k$ GCN layers and fine-tuning the rest of the network |
| GCN $k$ | Copying and fine-tuning the weights upto $k$ GCN layers from the base model while training the rest of the network from random initialization |
| Unfrozen | Fine-tuning the weights of all GCN layers after feature transfer |

## 3 Results and Discussion

We started with training the base models for brain decoding using task-fMRI data from HCP 1000 subjects. Using a 10s window with a time step of 3 TRs (equivalent TR=2.1s), the 21 cognitive states across six cognitive domains were identified with an average test accuracy of 84.17%. Using a 20s window, the decoding accuracy increased up to 89.36%. Higher accuracy was achieved in the domain-specific base models, for instance, distinguishing the five types of body movements (Motor task) with an accuracy of 93.48% using 5 fMRI volumes and classifying the eight types of visual working-memory tasks with an accuracy of 89.93% using 10 fMRI volumes.

### 3.1 Transfer Learning on datasets with the same scanning environment

In order to investigate the sample size effect on transfer learning, we generated three subsets from the HCP database consisting of 12, 24, and 36 subjects, all of which have not been used to train the base model. Due to the small sample size used in this analysis, all models were evaluated using a 10-fold cross-validation.

#### 3.1.1 Feature transfer performed better without fine-tuning

We observed a significant effect of sample size, not only on the model trained from scratch but also via feature transfer and fine-tuning (see Figure 1). First, the generalizability of the scratch model gradually improved as the available data samples enriched, for instance, 59%, 79% and 89% respectively for 12, 24 and 36 subjects when distinguishing the five types of body movements. Second, feature transfer largely compensated for the loss in generalizability due to the small sample size. With the aid of transfer learning, we achieved an improvement in decoding accuracy similar to what could be observed when doubling the sample size. For instance, the prediction accuracy on the Motor task increased up to 78%, 90% and 93% for 12, 24 and 36 subjects after feature transfer. Third, no further improvement was observed after fine-tuning the GCN layers even using a layer-wise approach (around 3% decrease). Similar findings were observed in the classification of working-memory tasks, with the decoding accuracy largely improved by increasing the sample size, for instance, 55%, 69% and 80% respectively for 12, 24 and 36 subjects when training the model from random initialization. The generalizability of brain decoding was highly improved by using features transferred from the base model, with the average gain as 16% for 12 subjects, 10% for 24 subjects, 2% for 36 subjects, but was weakened after fine-tuning the GCN layers with a decrease of around 4%.

**Figure 1:**
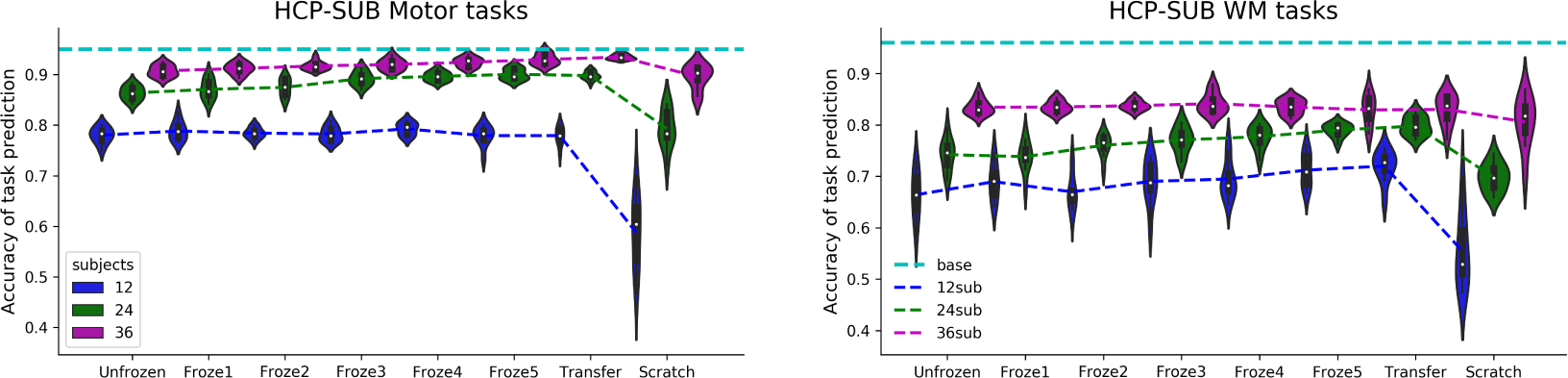
Transfer learning boost the decoding performance on the Motor (left) and Working-memory tasks (right) from the HCP subsets.

This is probably due to the fact that the neuroimaging data acquired from the same scanner using the same BOLD sequence and experimental protocols, can be embedded in the same low-dimensional manifold in which different task conditions can be easily separated. Thus, the representations transferred from the base model, that have been trained in a large population, provided an unbiased projection onto the manifold regardless of the small sample size and consequently improved the accuracy of task prediction. By contrast, fine-tuning the GCN layers forced the decoding model to adapt to the data distribution of the small training set and possibly caused overfitting. In order to verify this hypothesis, we performed additional analysis on the learned graph representations for different types of decoding models, including similarity analysis of learned representations, the nonlinear projections, as well as saliency maps.

#### 3.1.2 Similarity analysis of graph representations

In order to understand the benefits of transfer learning, we compared the learned representations between different layers and decoding models. The similarity of representations was calculated using centered kernel alignment (CKA) with a linear kernel which has been shown to be robust to different initializations (5). In the following similarity analysis of representations, we used the trained model resulting from the first cross-validation. Layer activation of the base model was first calculated on the HCP subsets of 24 subjects and then compared with other the decoding models either trained from random initialization, or using feature transfer with and without further fine-tuning. It is worth noting that the layer activation should be identical for the two models, since the transfer model used fixed representations extracted from the base model. Consistent with this theory, we found a block structure in the similarity matrix with CKA=1.0 in the diagonal. The similarity between GCN layers indicated that, different representations were learned for the 1st, 5th and 6th GCN layers (CKA<0.5), while highly similar representations were captured in the middle layers (2-4 GCN layers, mean CKA=0.95). The scratch model also showed a similar pattern (Figure 2A) with high similarities for the 1st, 5th and 6th GCN layers (mean CKA=0.85 comparing with the base model) but much lower similarities for the middle layers (mean CKA=0.68). The results indicated that the scratch model captured similar low- and high-level representations by training from random initialization on the small dataset, but learned very different representations for the middle layers. By contrast, the fine-tuning models retained the transferred representations to a large degree, with only small changes in the last GCN layers by gradually embedding more mid-layer representations in order to adapt to the specific dataset or population (mean CKA increasing from 0.47 to 0.77). Similar findings were observed for the working-memory tasks (see Section 3 and Figure S4 in Supplementary).

**Figure 2:**
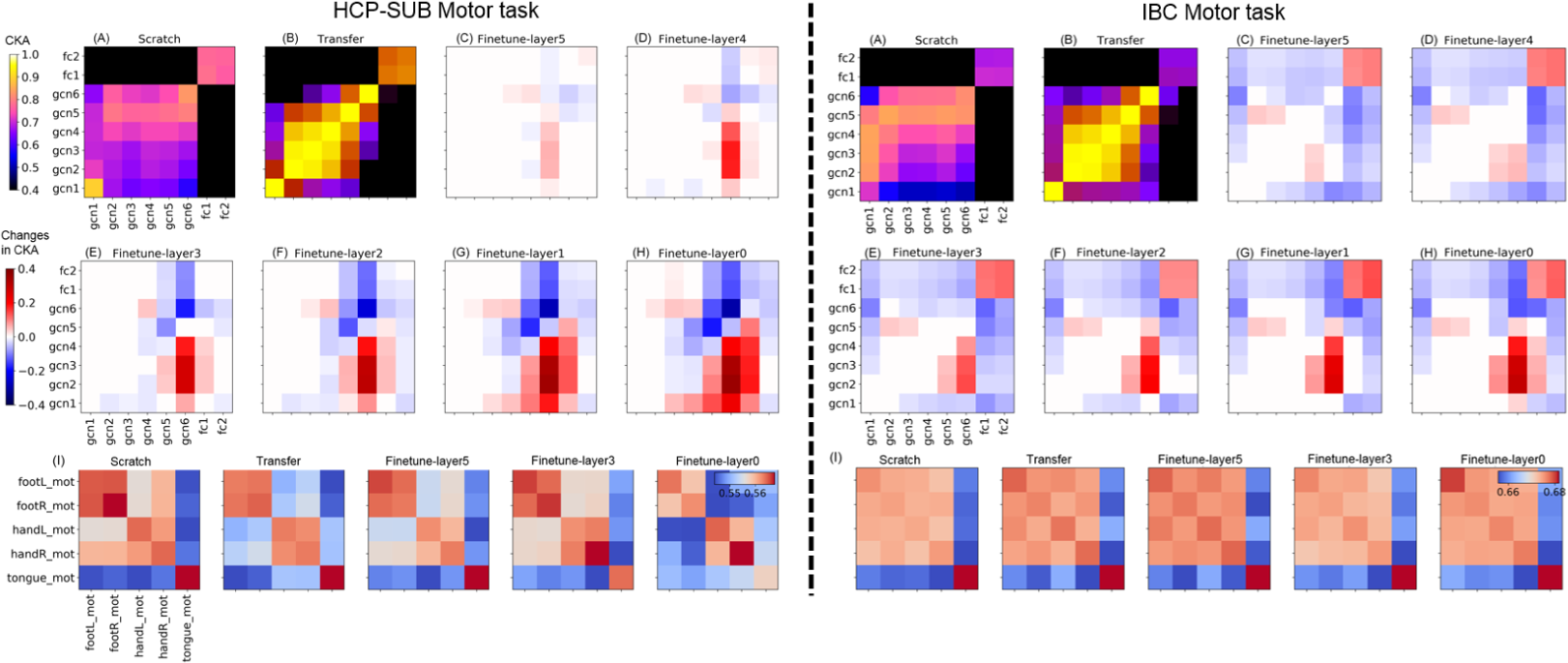
Similarity of representations for the decoding models after transfer learning and fine-tuning on the HCP-SUB (left) and IBC (right) Motor task. The learned representations of the decoding models were evaluated by comparing them with the base model using CKA. Eight different decoding models were evaluated, including training from scratch (A), using transferred graph representations (B) and fine-tuning GCN layers (C-H). (A-B) showed the raw CKA values between the chosen decoding model (x-axis) and the base model (y-axis), while (C-H) showed the changes in CKA values when comparing to the transfer model. The similarity of representations between tasks were calculated for each model (I) using the Mahalanobis distance.

#### 3.1.3 Projections of graph representations

The learned graph representations of brain dynamics were projected onto a 2-dimensional space using t-SNE (6). The similarity of representations between tasks were calculated using Mahalanobis distance. The transfer model achieved the highest decoding accuracy on the Motor task (92.5%) where the five types of body movements can be visually separated after the projection of learned graph representations (Figure 2I). Small mixing effect was still observed between the left- and right-side of body movements, for instance, between left and right-hand movement, which can be further distinguished when projecting the representations to a higher dimension (see Figure S6 in Supplementary). The scratch model achieved a much lower decoding accuracy (83.7%) with stronger mixing effects among the projections of left and right foot movement as well as right hand movement that were not separable even in a higher dimension. The decodability of the fine-tuning models gradually decreased after layer-wise fine-tuning (0.925, 0.9125, 0.8875 respectively for only tuning the last GCN layers, the last three GCN layers and all GCN layers). But the representations of different movements were gradually separated from each other, especially between left and right foot movements, as well as between left and right hand, with a tradeoff of smaller distinctions with the tongue movement. For the data distribution after projecting graph representations using t-SNE and other methods, see Figure S7 for Motor task and Figure S9 for Working-memory tasks in Supplementary.

#### 3.1.4 Saliency maps: brain activations and inter-subject variability

In order to investigate whether the decoding models learned a set of biologically meaningful features, we generated saliency maps on the trained model by propagating the non-negative gradients backwards to the input layer (12). An input feature is salient or important only if its little variation causes big changes in the decoding output. Thus, high values in the saliency map indicate large contributions during the prediction of task states. The saliency maps on the Motor task showed that different types of movements were associated with high salience in the primary motor and somatosensory cortices (Figure 3), which have been shown to be the main territories engaged during movements of the human body in the neuroscience literature (8). No clear somatotopic organization was identified here, which was somewhat expected because the primary motor and somatosensory cortex were parcellated into single strips in the Glasser’s atlas (Glasser et al. 2016). To compensate for this effect, other regions beyond the sensorimotor cortex were detected in the saliency map. For instance, regions around the operculum were selectively activated for the tongue movement (regions c and d in Figure 3), while BA2 was selectively activated for hand movement (region b in Figure 3). High consistency and notable divergence in the saliency maps were observed between different decoding models. First, a similar set of brain regions were detected across all decoding models, using biologically meaningful features during task prediction. Second, different brain signatures were generated from the neural dynamics of these brain regions. Specifically, the feature transfer model learned highly stable and task-distinctive brain signatures, consisting of regions selectively responding to foot (region a), hand (region b), and tongue movement (regions c and d), while suppressing their response to other movements. Fine-tuning the GCN layers slightly disrupted such clean and stable distinctiveness between tasks, for instance reduced response to foot movement in region a. High inter-subject variability was also detected in the saliency map which may infer the complexity of the trained model and be used as an indicator of model overfitting. When training the model from random initialization, we observed much higher inter-subject variability in the saliency maps, along with a random or high-order relationship between tasks and regions, which implied a high overfitting effect during model training by treating each subject differently.

**Figure 3:**
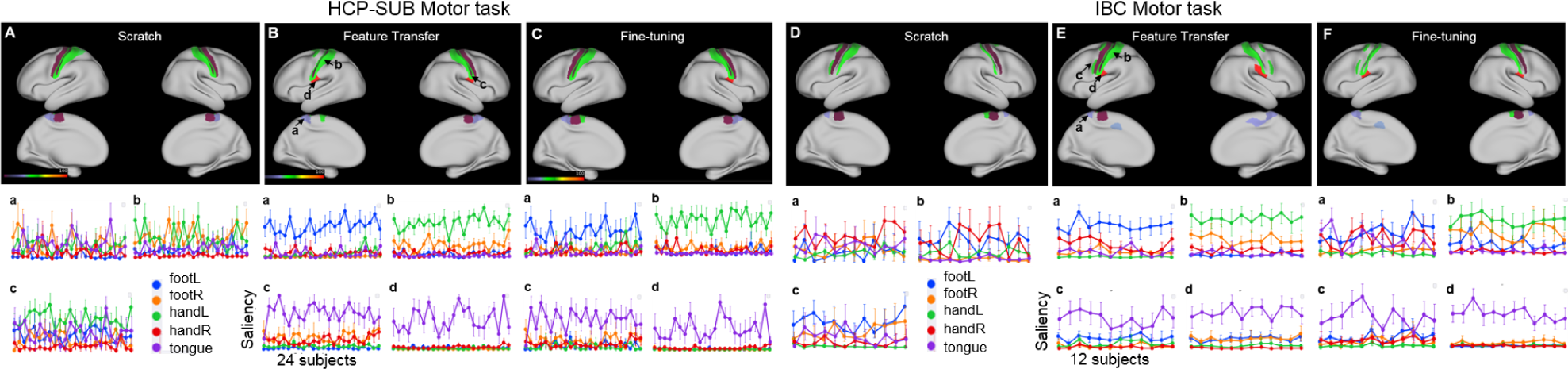
Saliency maps of decoding brain activities on the HCP-SUB (A-C) and IBC (D-F) Motor task. The effect of task conditions on the saliency maps were evaluated using repeated-measure ANOVA with “subject” as the random factor and “task” as the within-group factor. We only showed brain regions with a high saliency (>0.2) and having a significant effect of task (FWE corrected p<0.001) in the saliency maps. For Motor task, the data samples includes five types of movements, i.e. the movement of right foot (in blue), left foot (in orange), right hand (in green), left hand (in red), and tongue (in purple). The inter-subject variability was shown for four regions, including regions selectively responding to foot (region a), hand (region b), and tongue movement (regions c and d).

#### 3.1.5 Domain-specific transfer learning vs domain-general transfer learning

The above transfer-learning analysis was performed by using the domain-specific base model, for instance, the base model for the Motor task was trained only using fMRI data from the task sessions when participants performed the five types of body movements. An alternative approach was to train a domain-general base model using fMRI data from all available task domains. Smaller gains were obtained by transferring features from other cognitive tasks instead of copying domain-specific features, for instance, performance boost on Motor task as 10% for 12 subjects, 4% for 24 subjects, 2% for 36 subjects. Further fine-tuning the GCN layers provided an additional boost of 1%. Another approach was to transfer only a part of graph representations instead of copying features of all GCN layers. For within-domain transfer learning, higher boost on decoding performance was achieved by transferring all GCN layers. On the contrary, for across-domain transfer learning, the best performance was achieved by only transferring and fine-tuning the first 4 GCN layers, with performance boost over the scratch model as 12% for 12 subjects, 6.8% for 24 subjects, 2% for 36 subjects. Our results indicated that the low-level representations of brain dynamics were transferable among different cognitive tasks, while both low and high-level representations were shared within the same task domain and transferable among different groups of subjects. In both cases, the benefits of transfer learning were dismissed as the sample size of the training set increased.

### 3.2 Transfer Learning on datasets with different scanning conditions

We tested the above pipeline on an independent dataset consisting of 12 subjects from the Individual Brain Charting (IBC) database, where fMRI data were acquired using a different scanner, different BOLD sequences, and different repetition time (TR=2s). For the Motor task, training from random initialization using less than 1k data samples resulted in an effective decoding (average test accuracy=91% across 10-fold cross-validation, random chance=20%). Still, using features transferred from the domain-specific base model provided an additional gain of 7%, but with no further improvement or even slightly lower performance after fine-tuning the GCN layers. Interestingly, the direct prediction on the IBC Motor task using the base model on HCP subjects yielded a very high decoding accuracy (97.5%). These results indicated that the neural representations of the Motor task were highly stable not only between subjects but also across sites. Further fine-tuning learned site-specific features but may overfit on the small dataset (Figure 4). We found consistent but somewhat distinct patterns on the Working-memory task. First, the base model trained on HCP subjects showed poor generalizability on IBC subjects (test accuracy=30%, chance level=12.5%), coinciding with the literature that high inter-subject variability has been reported in both behavior and neural activity of the Working-memory tasks (2; 7). On the contrary, the decoding model trained from random initialization performed much better (41%), with a performance boost of 5% by using features transferred from the base model and an additional gain of 4% after further fine-tuning the GCN layers. Our results indicated a strong site effect in the learned representations of brain dynamics for the Working-memory task, but with a smaller effect of task domain (see Table 2).

**Table 2:** Performance boost of transfer learning on IBC Motor and Working-memory tasks

| Models | ALLTask-Motor |  | ALLTask-WM |  | MOTOR-Motor |  |  | WM-WM |  |  |
| --- | --- | --- | --- | --- | --- | --- | --- | --- | --- | --- |
|  | Transfer-Learning (%) |  | Transfer-Learning (%) |  | Transfer-Learning (%) |  |  | Transfer-Learning (%) |  |  |
|  | Boost<br>over<br>Scratch<br>(90.3) | Boost<br>over<br>Transfer<br>(93.8) | Boost<br>over<br>Scratch<br>(43.7) | Boost<br>over<br>Transfer<br>(43.8) | Boost<br>over<br>Scratch<br>(89.5) | Boost<br>over<br>Base<br>(97.5) | Boost<br>over<br>Transfer<br>(96.7) | Boost<br>over<br>Scratch<br>(42) | Boost<br>over<br>Base<br>(30) | Boost<br>over<br>Transfer<br>(47) |
| GCN 1 | 2.3 | -0.5 | 7.1 | 4.5 | 2.2 | -4.1 | -5 | 4 | 15 | -1 |
| GCN 2 | 2.4 | 0.7 | 7.2 | 4.6 | 1.9 | -4.4 | -5.3 | 6.2 | 17 | 1.2 |
| **GCN 3** | **3.7** | **1.0** | **8.6** | **6.0** | 2.9 | -3.4 | -4.3 | 7.6 | 18.8 | 2.7 |
| GCN 4 | 3.1 | 0.4 | 8.2 | 5.6 | 2.9 | -3.4 | -4.3 | 7.5 | 18.7 | 2.6 |
| GCN 5 | 2.4 | 0 | 5.6 | 3 | 5 | -2.1 | -3 | 8.3 | 19.4 | 3.3 |
| **GCN 6** | 2.3 | 0 | 0.4 | -2 | **7.2** | **-1.3** | **-2.2** | **9** | **21** | **4** |

**Figure 4:**
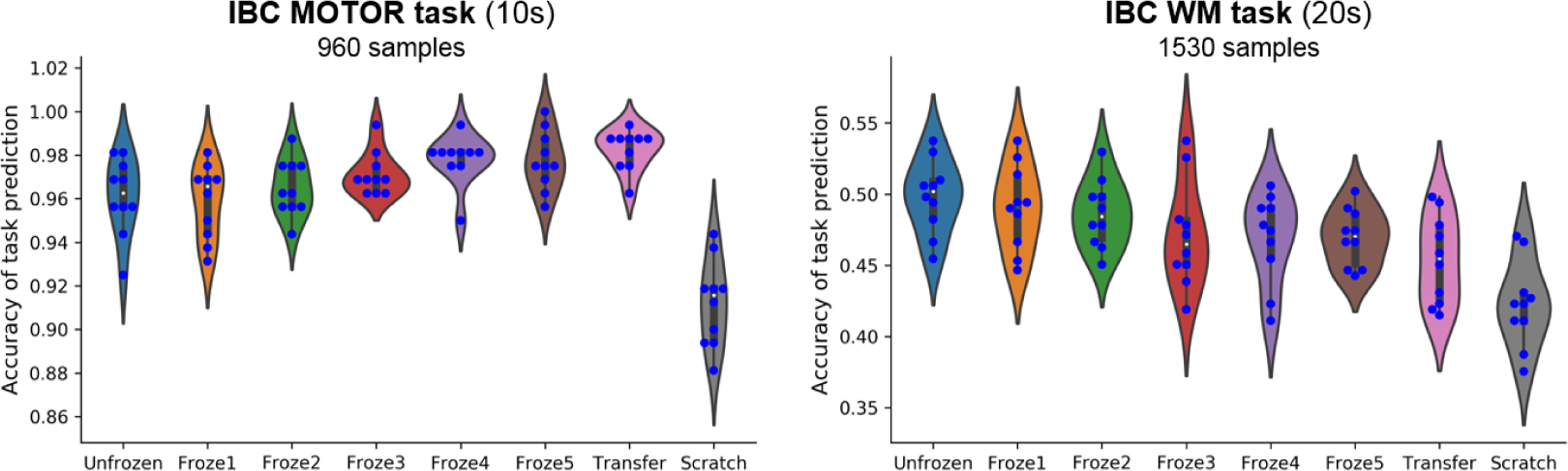
Transfer learning boosts the decoding performance on the Motor (left) and Working-memory (right) tasks from the IBC database

These observations somewhat coincided with the findings of transfer learning on natural images that features from the first few layers were generalized across different tasks while features from higher layers were more specific to a particular task (15). In our experiments, we also found that the representations of brain activities from the first three GCN layers showed high generalizability across different task domains, while the high-level representations (from 5th and 6th GCN layers) were only generalizable within the same task domain especially for fMRI data acquired from a different scanning environment. On the other hand, unlike natural images, the scanning conditions had a much higher impact on the transferability of graph representations of brain dynamics as, for instance, the performance boost was reduced from 20% to 7% for the Motor task, and from 15% to 10% for the Working-memory tasks when transferring across different sites.

#### 3.2.1 Similarity of graph representations

The scratch model captured similar representations as the base model for high-level GCN layers (mean CKA=0.82), as shown in Figure 2I. By contrast, the first GCN layer embedded all mid-layer representations of the base model (mean CKA=0.84), while retaining a much lower similarity with the low-level representations (CKA=0.71), which may indicate an overfitting effect in the scratch model by learning complex representations in the low-level GCN layers. Using a layer-wise fine-tuning approach after feature transfer, the last GCN layers gradually enhanced the embedding of the midlayer representations of the base model (mean CKA increasing from 0.50 to 0.78), while retaining the low-level representations to a large degree. Similar findings were observed when projecting the graph representations onto 2-dimensional spaces, for instance, fine-tuning GCN layers gradually separated the representations for the left- and right-side of body movements while learned highly similar representations for movements of the same category (see Figure S8 in Supplementary).

#### 3.2.2 Saliency maps: brain activations and inter-subject variability

We compared the saliency maps of different decoding models on the IBC Motor fMRI dataset (see Figure 3). A similar set of brain regions were detected that were highly contributed to the classification of different types of movements, consisting of the sensorimotor cortex and regions around the operculum. The transferred model learned stable and task-distinctive brain signatures by including regions selectively responding to foot (region a), hand (region b), and tongue movement (regions c and d). After fine-tuning the GCN layers, the model learned more complex representations along with high inter-subject variability in the saliency maps, especially for the hand and foot movements in regions a and b. When training the model from random initialization, a much higher inter-subject variability was detected, along with complex combinations among different movements, which implied a strong overfitting effect on the small dataset by learning different representations on each individual and using subject-specific decoding inference.

## 4 Conclusion

We proposed a new pipeline for transfer learning on medical imaging that used a base model trained on a large-scale neuroimaging dataset instead of transferring features from natural images. We evaluated the transferability of brain decoding across different cognitive domains, different groups of subjects, and even between different sites using different scanning sequences. Our results suggested that the learned representations of brain dynamics were highly transferable not only within the same cognitive domain but also across different task domains, as well as to new datasets acquired from a different scanning environment. Different from transfer learning on natural images, where the similarity of task domains played an important role (15), we observed a small effect in the domain of the base model, yielding a significant performance boost for both within-domain and across-domain transfer learning. By contrast, the scanning condition showed a much higher impact on the transferability of representations. Fine-tuning can further boost the decoding performance by learning more site-specific high-level features while retaining the transferred low-level representations of brain dynamics. Our study suggests a great potential of transfer learning and domain adaptation in medical imaging, possibly making contributions in a variety of domains, including neurological and psychiatric disorders.

