## Supplementary figures and images for "Transferability of Brain decoding using Graph Convolutional Networks"

### Supplement Figure 1

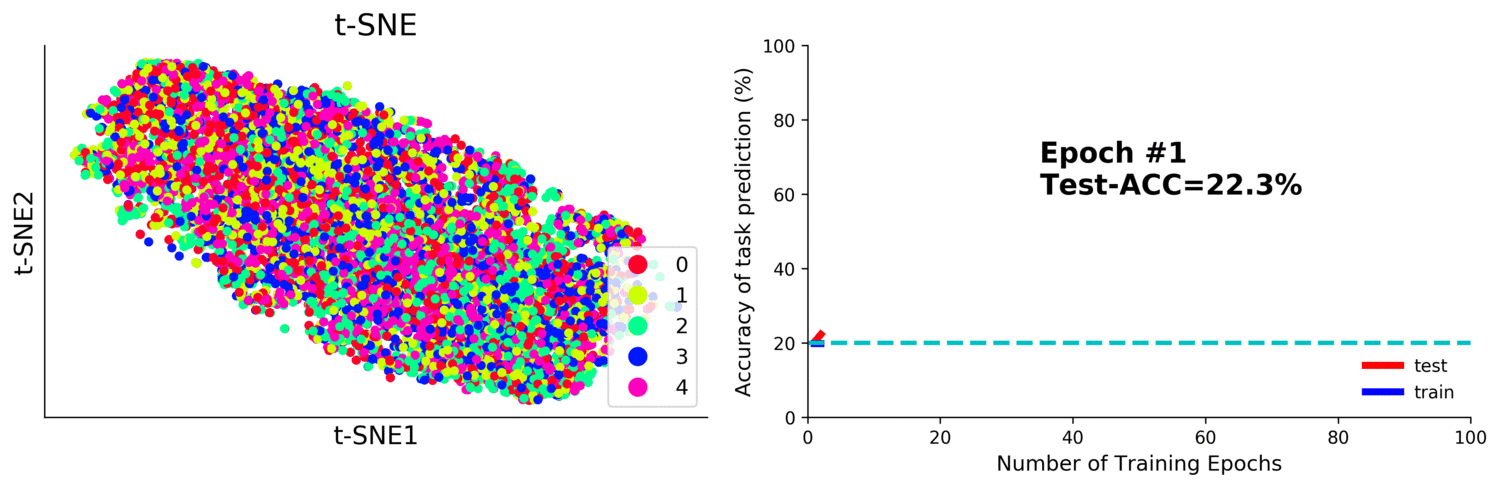

### Supplement Figure 2

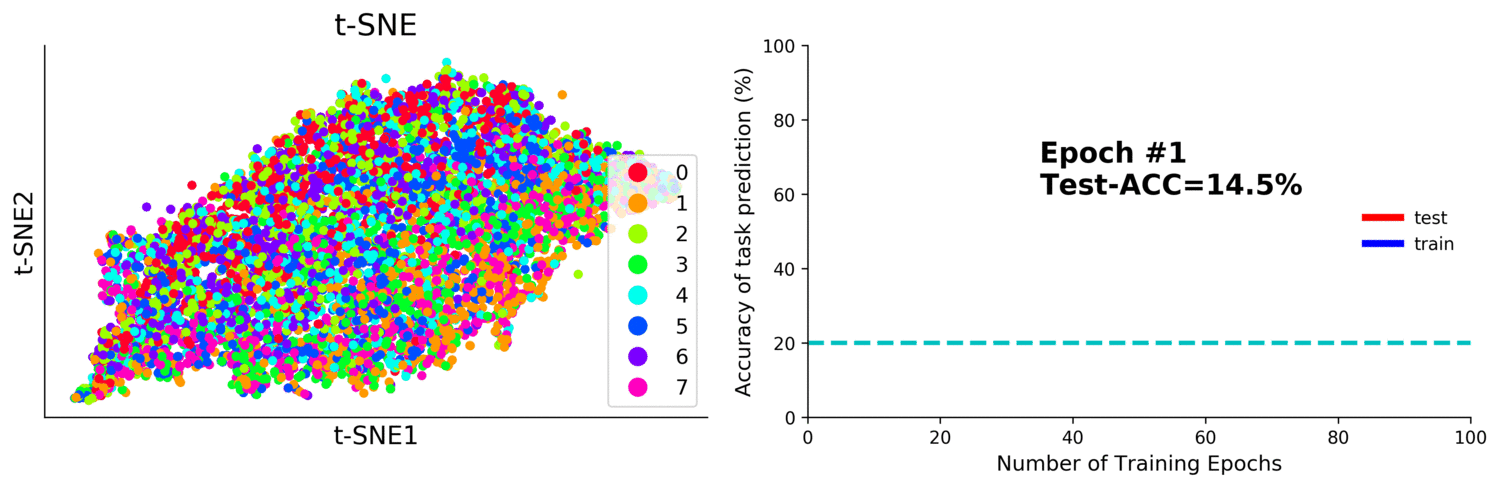
