## Supplement Materials for "Transferability of Brain decoding using Graph Convolutional Networks"

### Supplementary Materials

1. Visualization the train curve and learned representations for the base model

**Figure S1. Visualization of learned representations of Motor task-fMRI data during the training process of the base model.**
The layer activations of the last graph convolutional layer of the decoding model were extracted as the graph representations of brain dynamics. The learned representations were then projected onto a 2-dimensional space using t-SNE (left panel) (Maaten and Hinton 2008). The data samples includes five types of movements, i.e. the movement of right foot (class 0, in red), left foot (class 1, in cyan), right hand (class 2, in green), left hand (class 3, in blue), and tongue (class 4, in purple). The decoding accuracies on the training (blue line) and test (red line) sets were also collected during the model training process and shown in the right panel. The chance level of classifying the five types of movements was 20%, marked as a cyan line in the plot. Note that, the decoding performance started at the chance level of 20% and quickly reached the plateau of 97% after 20 training epochs.

**Figure S2. Visualization of learned representations of Working-memory task-fMRI data during the training process of the base model.**
The graph representations of brain dynamics were then projected onto a 2-dimensional space using t-SNE (left panel) (Maaten and Hinton 2008). The data samples includes eight types of visual working-memory, i.e. the recognition of images of places (class 2, in green), tools (class 2, in cyan), faces (class 1, in orange) and body parts (class 0, in red) and their counterparts for 2-back working memory tasks (classes 5 to 8). The decoding accuracies on the training (blue line) and test (red line) sets were also collected during the model training process and shown in the right panel. The chance level of classifying the five types of movements was 12.5%, marked as a cyan line in the plot. Note that, the decoding performance started at the chance level of 12.5% and quickly reached the plateau of 90% after 20 training epochs.

2. Transfusion by only copying upto ***k*** GCN layers

We also evaluated a different type of transfer learning named transfusion (Raghu et al. 2019), which only transfers the first few GCN layers while training the rest of the network from random initialization. For within-domain transfer learning, the best performance was achieved by transferring representations from all GCN layers. For across-domain transfer learning, the best performance was achieved only transferring and fine-tuning the first 4 GCN layers from the domain-general base model.

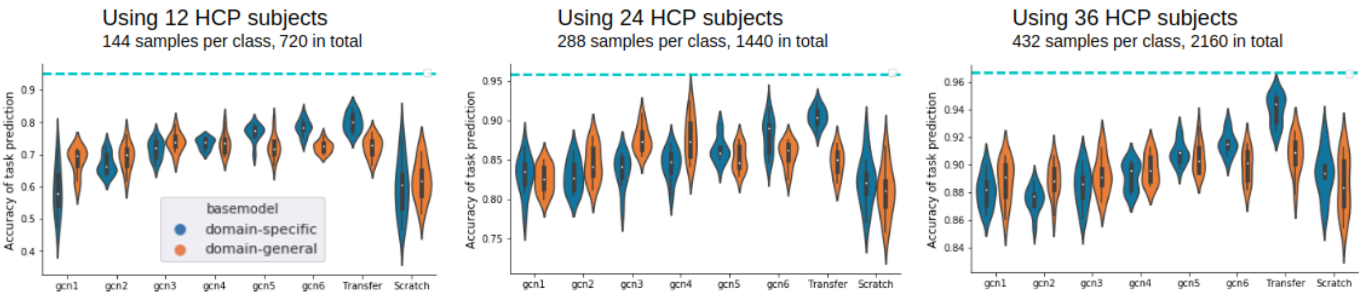


**Figure S3-1. Domain-specific transfer learning showed larger gains over domain-general transfer learning on the HCP SUB Motor task.**
We transferred the representations from upto k (ranging from 1 to 6) GCN layers from either domain-specific (e.g. Motor-Motor transfer learning, in blue) or domain-general (e.g. ALLTask-Motor transfer learning, in orange) base model and fine-tuned the entire decoding model. Different findings were shown for the two types of transfer learning. For within-domain transfer learning, transferring all learned representations from the base model achieved the best performance. For across-domain transfer learning, only using representations from the first four GCN layers provided the highest performance boost.

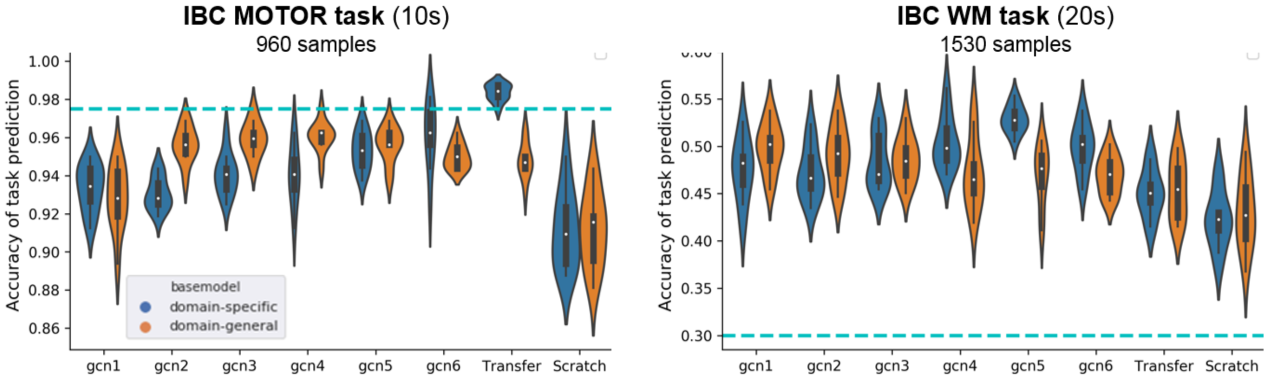


**Figure S3-2. Domain-specific and domain-general transfer learning on the IBC Motor (left) and Working-memory (right) tasks.**
Compared to within-domain transfer learning (in blue), the across-domain transfer learning (in orange) provided a smaller gain via fixed feature transfer, i.e. 2% for both Motor (left panel) and Working-memory tasks (right panel). For within-domain transfer learning, the best performance was achieved by transferring all learned representations from the base model. For across-domain transfer learning, the highest boost was achieved by copying and fine-tuning the first three GCN layers while training the rest of the network from random initialization (gain of 4% and 9% respectively for Motor and Working-memory tasks).

#### 3. Similarity of graph representations for Working-memory tasks

Layer activations of the different decoding models were compared with the base model using CKA. A block structure was detected for the between-layer similarity of the base model when comparing the layer activations between the transfer and the base model on the HCP SUB Working-memory tasks, which indicated that different representations were learned for the low and high GCN layers (CKA=0.5), while the middle layers learned a transition between the two blocks (CKA=0.88 and 0.87). The scratch model embedded the middle and high-level representations of the base model in the first GCN layer (CKA>0.84), but different from the low-level features, which may indicate a strong overfitting effect in the scratch model by learning highly complex representations in the early layers. The fine-tuning models retained the transferred representations to a large degree especially for the 1st and 2nd GCN layers, but with modifications in the last GCN layers which gradually embedded more low and mid-layer representations of the base model after layer-wise fine-tuning (mean CKA increasing from 0.55 to 0.85).

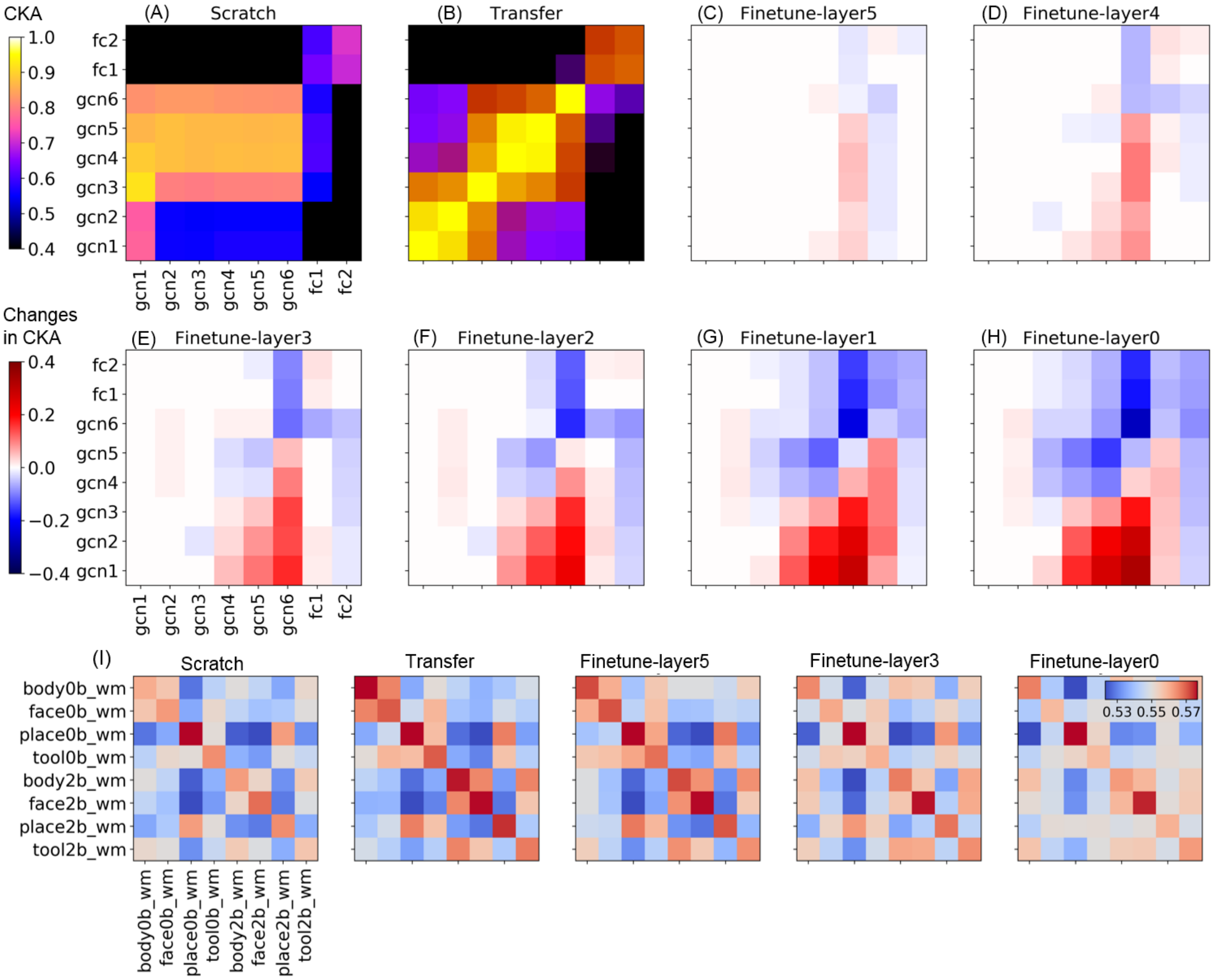


**Figure S4. Representational similarity of the decoding models after transfer learning and fine-tuning on the HCP SUB Working-memory task.**

The learned representations of the decoding models trained on HCP SUB Working-memory task, including 2304 samples of 20s fMRI time-series with TR=2.1s, were evaluated by comparing them with the base model using CKA. Eight different decoding models were evaluated, including training from scratch (A), using transferred graph representations (B) and fine-tuning GCN layers (C-H). (A-B) showed the raw CKA values between the decoding model (x-axis) and the base model (y-axis), while (C-H) showed the changes in CKA values when comparing to the transfer model. The similarity of representations between tasks were calculated for each model (I) using the Mahalanobis distance.

Similar findings were observed for the IBC Working-memory task. The same block structure was detected when comparing layer activations of the transfer and base model. Very different representations were learned when training the model from random initialization (Figure S5 A), except that the 1st GCN layer captured similar representations as the low and middle GCN layers from the base model (CKA=0.9 and 0.76 respectively). The fine-tuning models kept most of the low and middle-level representations from the base model, but gradually embedded more low-level features of the base model in the last GCN and 1st fully-connected (FC) layers after tuning until the 3rd GCN layer (mean CKA increasing from 0.52 to 0.76 for last GCN layer, from 0.31 to 0.58 for 1st FC layer).

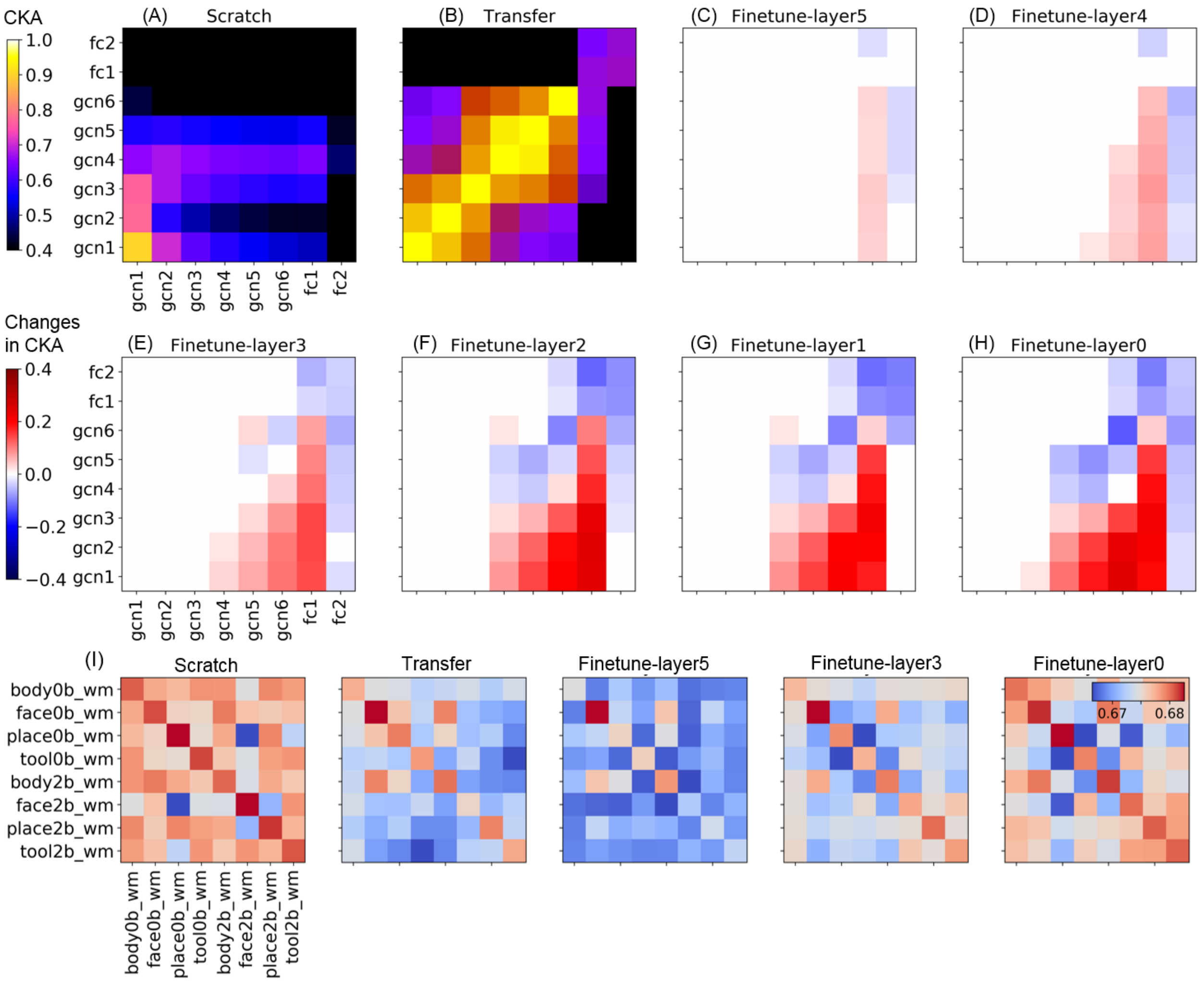

**Figure S5. Representational similarity of on the decoding model after transfer learning and fine-tuning on the IBC Working-memory task.**
The learned representations of the decoding models trained on HCP SUB Working-memory task, including 1530 samples of 20s fMRI time-series with TR=2s, were evaluated by comparing them with the base model using CKA. Eight different decoding models were evaluated, including training from scratch (A), using transferred graph representations (B) and fine-tuning GCN layers (C-H). (A-B) showed the raw CKA values between the decoding model (x-axis) and the base model (y-axis), while (C-H) showed the changes in CKA values when comparing to the transfer model. The similarity of representations between tasks were calculated for each model (I) using the Mahalanobis distance.

#### 4. Visualization of graph representations of brain dynamics for the Motor task

On the HCP SUB Motor task, despite the high decoding performance, we still observed a small mixing effect between the left- and right-side of body movements, for instance, between left and right-hand movement, which can be further distinguished when projecting the representations to a higher dimension (Figure S6). Similar to the HCP database, we found similar patterns in the IBC dataset (Figure S8): 1) the transfer model clearly differentiated the five types of body movements with the highest similarity to samples of the same category, but with minor mixing effect between foot and hand movements; 2) After layer-wise fine-tuning, the mixing effect was highly suppressed by showing an approximate diagonal matrix for between-class similarities; 3) the scratch only showed a clear distinction between tongue movement and other types of movements, while highly mixed between foot and hand movements. These different patterns of representational similarities can be used to explain their differences in the decoding performance.
It’s worth noting that, for the analysis on graph representations, we only used the trained model resulting from the 1st cross-validation, while the results reported in the main text were averaged 10-fold cross-validations. Thus the accuracy reported here might be different from the main text, mainly due to the random effect in 1) different splits of dataset; 2) different initializations for model training.

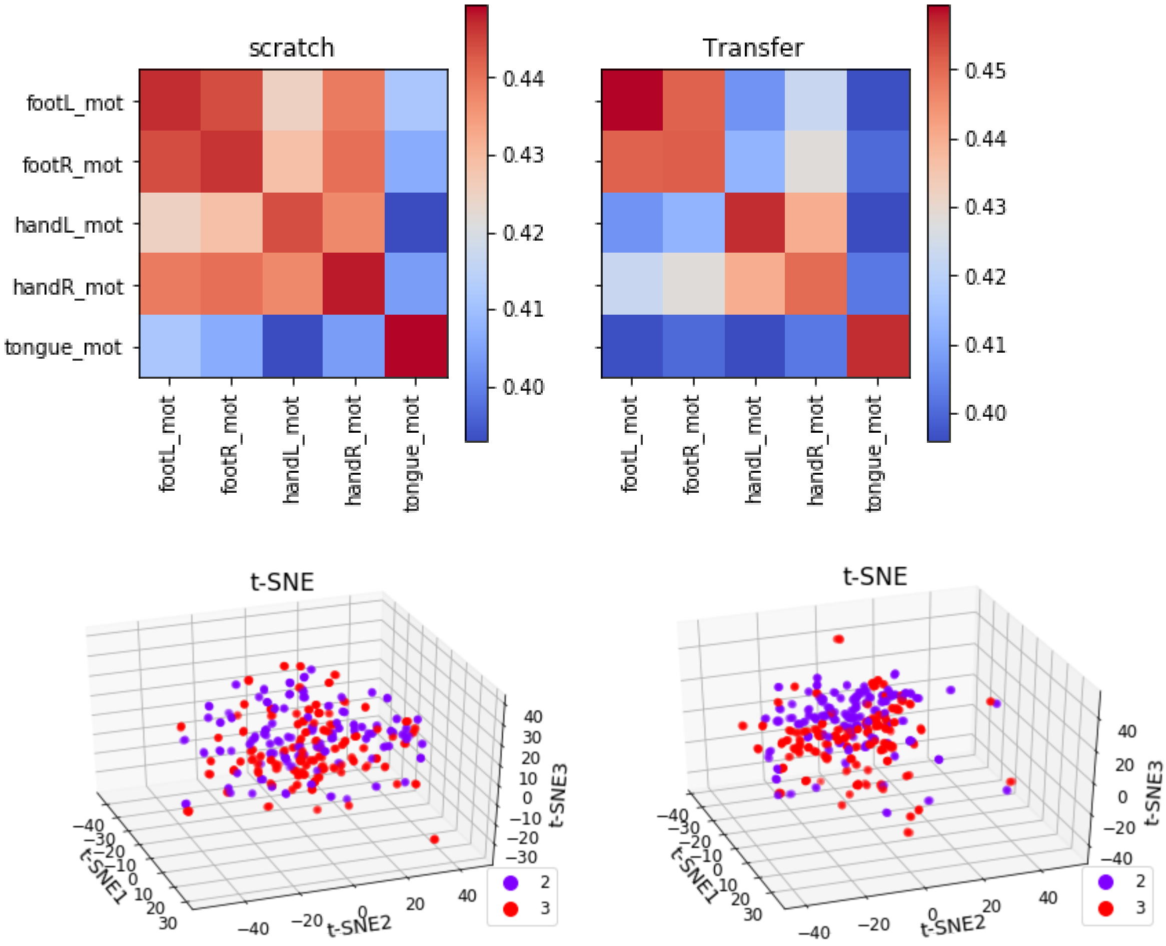


**Figure S6. Projecting the representations of left and right hand movements onto a 3-dimensional space on the HCP SUB Motor task.**

The representations of left and right hand movements were highly overlapped in the scratch model (left panel), along with a mixing effect between the movement of right hand and foot as well. By contrast, in the transfer model (right panel), the left (in purple) and right hand (in red) movements were separable using a separating hyperplane, with additional suppression on the similarity between hand and foot movements.

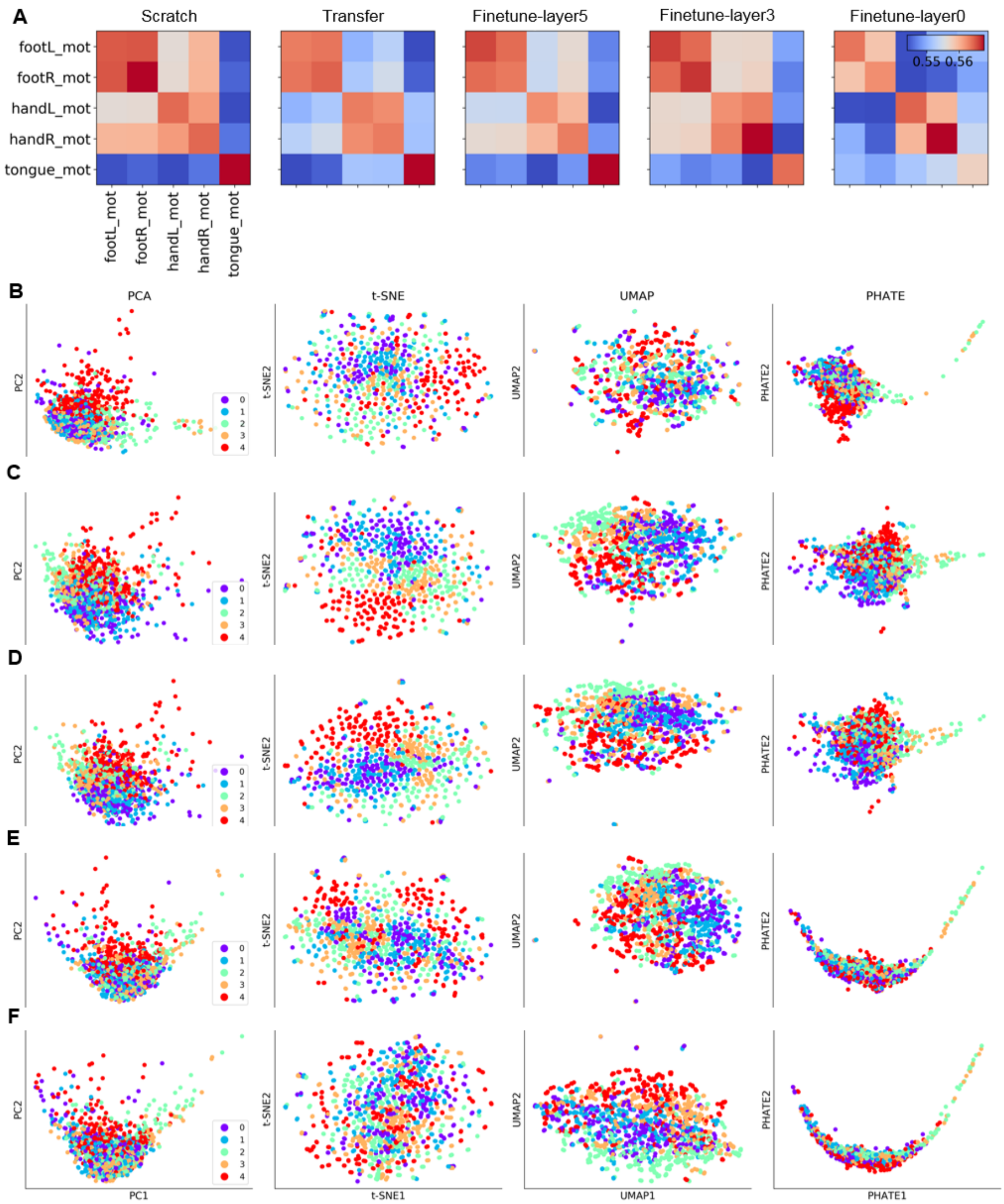


**Figure S7. Visualization of learned graph representations for transfer learning and fine-tuning on the HCP SUB Motor task.**

The decoding models were trained on HCP SUB Motor task, including 1440 samples of 10s fMRI time-series with TR=2.1s. We then projected the learned representations of all data samples onto a 2-dimensional space by using different dimension reduction techniques, including PCA (1st column), t-SNE (2nd column), UMAP (3rd column) and PHATE (last column). For Motor task, the data samples includes five types of movements, i.e. the movement of right foot (class 0, in purple), left foot (class 1, in green), right hand (class 2, in cyan), left hand (class 3, in orange), and tongue (class 4, in red). The similarity of representations between different types of movements were calculated for the decoding models trained from scratch, using features transfer with and without fine-tuning (A). Among these models, the transfer model achieved the highest decoding accuracy (C: 92.5%) and gradually decreased after layer-wise fine-tuning ( 92.5%, 91.2%, 88.7%

respectively for fine-tuning the last 1 (D),3 (E), and all GCN layers (F)). Lowest accuracy was achieved by training the model from scratch (B: 83.7%).

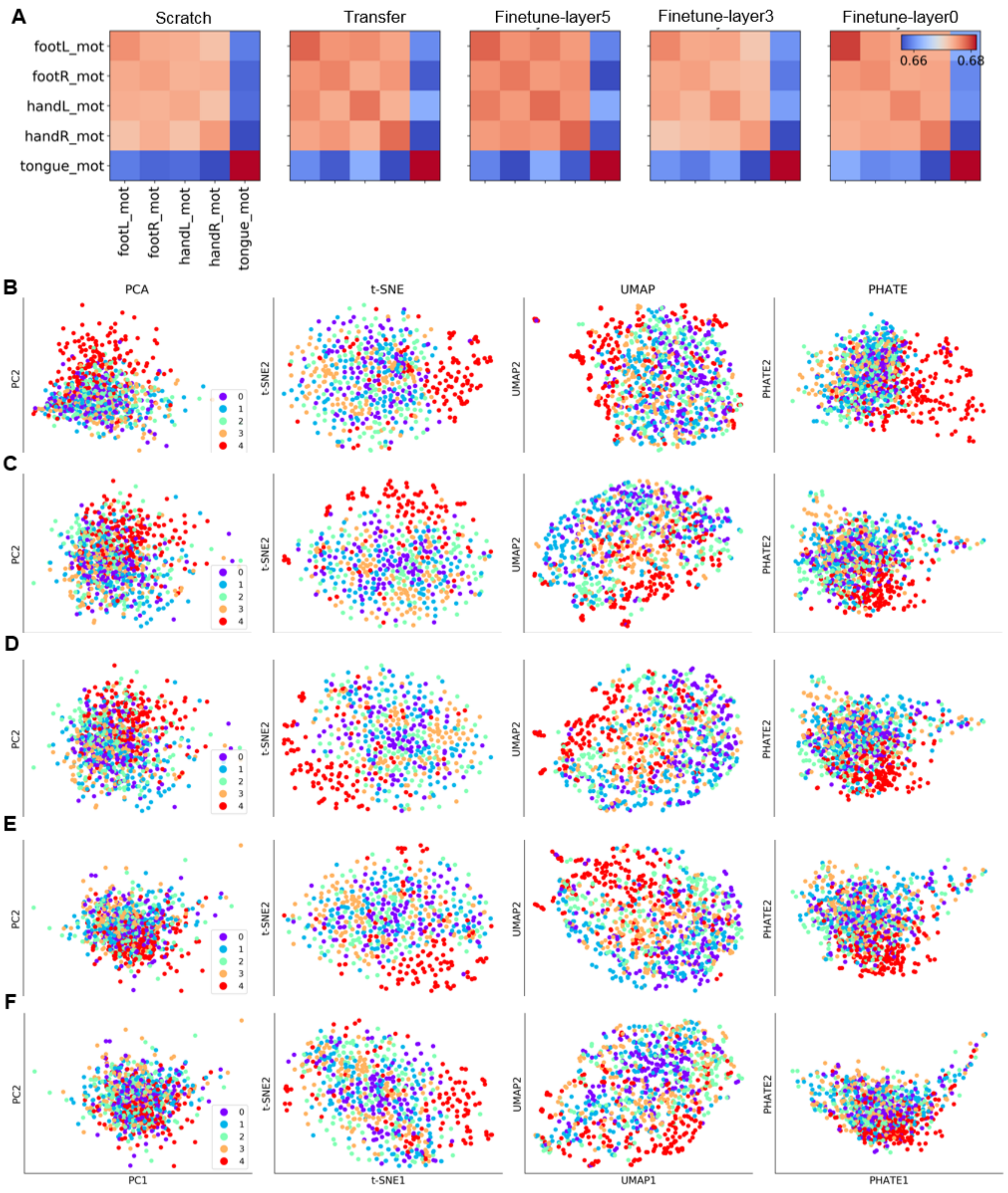


**Figure S8. Visualization of learned graph representations for transfer learning and fine-tuning on the IBC Motor task.**

The decoding models were trained on IBC Motor task, including 960 samples of 10s fMRI time-series with TR=2s. We then projected the learned representations of all data samples onto a 2-dimensional space by using different dimension reduction techniques, including PCA (1st column), t-SNE (2nd column), UMAP (3rd column) and PHATE (last column). For Motor task, the data samples includes five types of movements, i.e. the movement of right foot (class 0, in purple), left foot (class 1, in green), right hand (class 2, in cyan), left hand (class 3, in orange), and tongue (class 4, in red). The similarity of representations between different types of movements were calculated for the decoding models trained from scratch, using features transfer with and without fine-tuning (A). High decoding accuracy was achieved on the transfer model (C: 96.88%), which slightly decreased after layer-wise fine-tuning (95.63%, 95.63% and 96.25% respectively for fine-tuning the last 1 (D),3 (E), and all GCN layers (F)). Lowest accuracy was achieved by training the model from scratch (B: 90.62%).

#### 5. Visualization of graph representations of brain dynamics for Working-memory tasks

We also projected the learned graph representations of brain dynamics during the Working-memory task onto 2-dimensional spaces. On the HCP SUB Working-memory tasks, the scratch model showed highly distinctive representations of the eight types of visual working-memory tasks (Figure S9). Yet, small mixing effects were present, for instance, between recognition of face and body images, and between 0-back and 2-back recognition of place images. The segregation of tasks was highly strengthened in the feature transfer model by largely increasing the similarity within each category. However, moderate similarity between different tasks was still shown in the transfer model.  After fine-tuning the GCN layers, the between-class similarities were gradually suppressed, but come with a price of reducing within-class similarities as well. As a result, the decoding accuracy did not improve but decreased after fine-tuning. Different effect was found on the IBC Working-memory tasks (Figure S10). After projecting the learned graph representations of brain dynamics onto 2-dimensional spaces, we found that the representations learned from scratch were highly randomized with a strong mixing effect between different types of visual tasks. The scratch model only identified a clear distinction between the recognition of face and place images. The transfer model showed an approximate diagonal for between-class representational similarities. But it also showed highly diverse representations for both 0-back and 2-back face recognition, for instance, samples of the 0-back face recognition task were mixed with 0-back recognition of place images and 2-back recognition of body images. After fine-tuning the GCN layers, the diversity within each category was highly suppressed  with the highest similarity to samples of the same category for all task conditions. But the mixing effect was still present, for instance, moderately similar representations between 0-back face recognition and 2-back recognition of body images. These results implied that more data were required in order to further suppress the inter-class similarity in representations.

It’s worth noting that, for the analysis on graph representations, we only used the trained model resulting from the 1st cross-validation, while the results reported in the main text were averaged 10-fold cross-validations. Thus the accuracy reported here might be different from the main results.

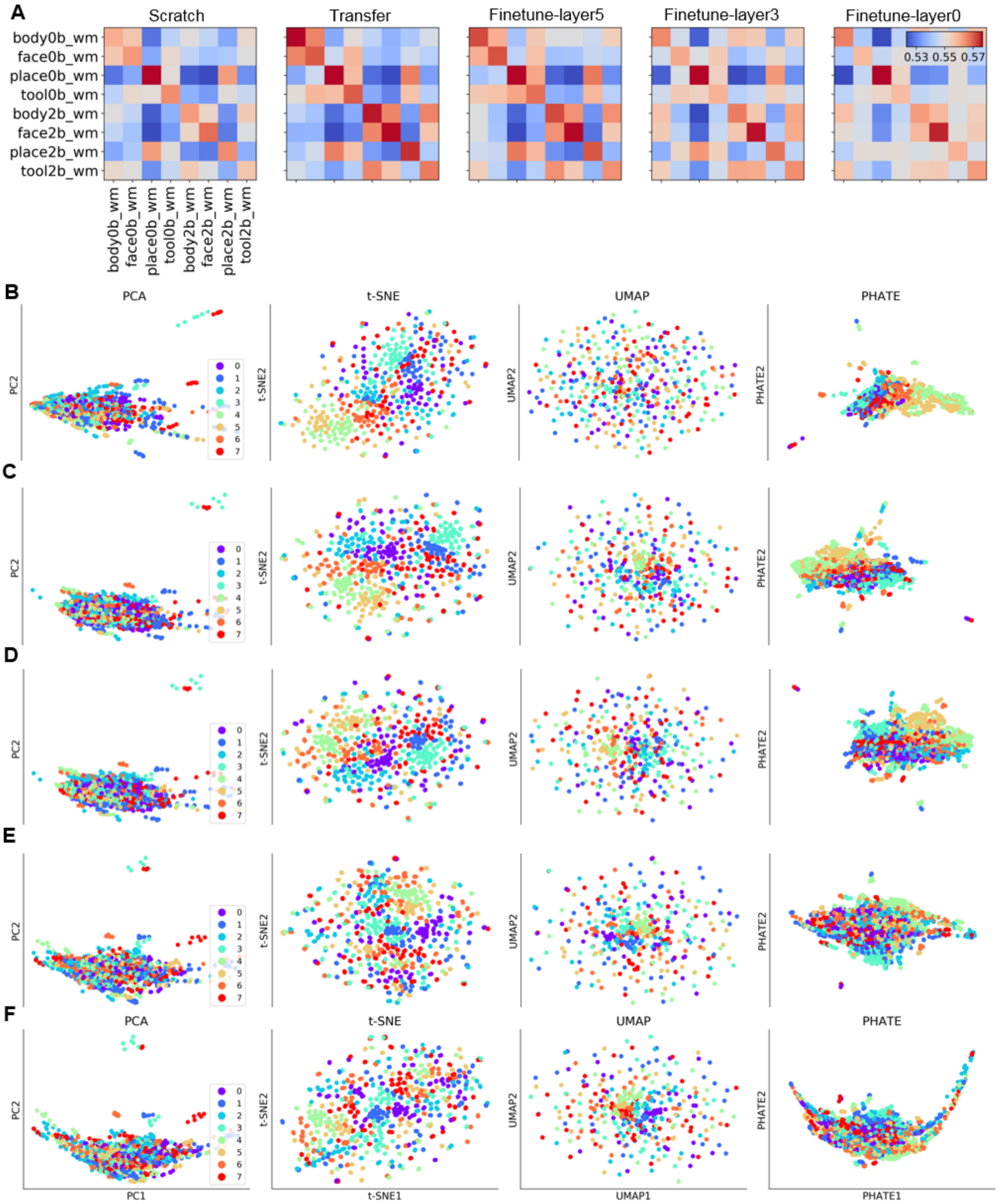

**Figure S9. Visualization of learned graph representations for transfer learning and fine-tuning on the HCP SUB Working-memory task.**

The decoding models were trained on HCP SUB Working-memory task, including 2304 samples of 20s fMRI time-series with TR=2.1s. We then projected the learned representations of all data samples onto a 2-dimensional space by using different dimension reduction techniques, including PCA (1st column), t-SNE (2nd column), UMAP (3rd column) and PHATE (last column).
For Motor task, the data samples includes five types of movements, i.e. the 0-back (class 0, in purple) and 2-back recognition of body (class 4, in light cyan), 0-back (class 1, in blue) and 2-back recognition of face (class 5, in light orange), 0-back (class 2, in green) and 2-back recognition of place (class 6, in orange), and 0-back (class 3, in cyan) and 2-back recognition of tool images (class 7, in red). The similarity of representations between different types of movements were

calculated for the decoding models trained from scratch, using features transfer with and without fine-tuning (A). Among these models, the transfer model achieved the second highest decoding accuracy (C: 79.1%) and slightly improved after fine-tuning the last GCN layer (D: 80.2%), and then gradually decreased after layer-wise fine-tuning (E: 78.9%, F: 75.7% respectively for fine-tuning the last 3 and all GCN layers). Lowest accuracy was achieved by training the model from scratch (B: 64.8%).

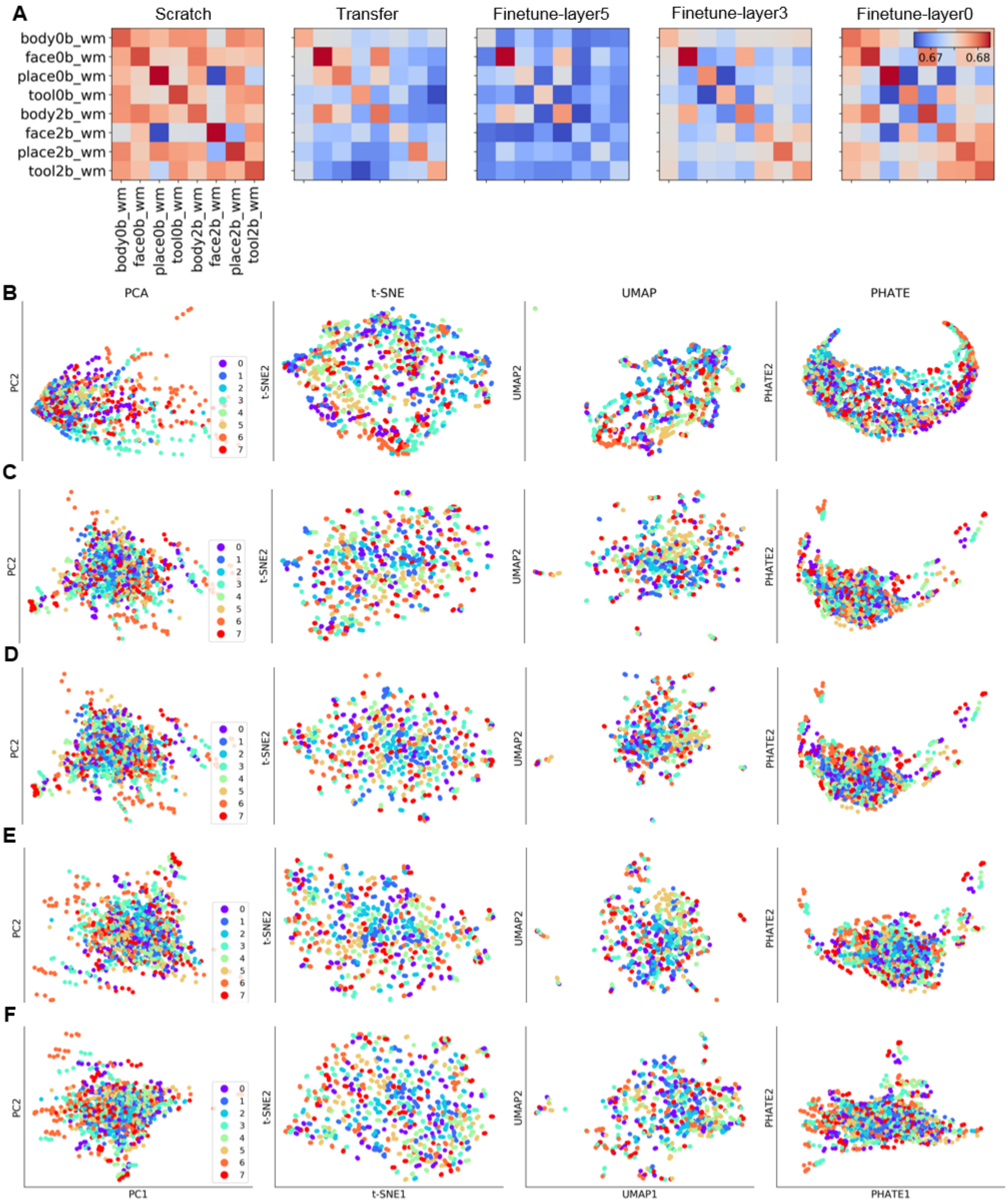


**Figure S10. Visualization of learned graph representations for transfer learning and fine-tuning on the IBC Working-memory task.**

The decoding models were trained on the IBC Working-memory tasks, including 1530 samples of 20s fMRI time-series with TR=2s. We then projected the learned representations of all data samples onto a 2-dimensional space by using different dimension reduction techniques, including PCA (1st column), t-SNE (2nd column), UMAP (3rd column) and PHATE (last column).
For Motor task, the data samples includes five types of movements, i.e. the 0-back (class 0, in purple) and 2-back recognition of body (class 4, in light cyan), 0-back (class 1, in blue) and 2-back recognition of face (class 5, in light orange), 0-back (class 2, in green) and 2-back recognition of place (class 6, in orange), and 0-back (class 3, in cyan) and 2-back recognition of tool images (class 7, in red). The similarity of representations between different types of movements were calculated for the decoding models trained from scratch, using features transfer with and without fine-tuning (A). The transfer model achieved a low decoding accuracy (C: 48.1%) which was further reduced after fine-tuning the last GCN layer (D: 47.8%), and then gradually increased to the top performance after layer-wise fine-tuning (E: 49.8%, F: 51.3% respectively for fine-tuning the last 3 and all GCN layers). A much lower accuracy was achieved by training the model from scratch (B: 39.9%).
